# The choice of the environmental covariate affects the power to detect variation in reaction norm slopes

**DOI:** 10.1101/311217

**Authors:** Phillip Gienapp

## Abstract

Many traits are phenotypically plastic, i.e. the same genotype expresses different phenotypes depending on the environment. Individuals and genotypes can vary in this response to the environment and this individual (IxE) and genetic variation in reaction norm slopes (GxE) can have important ecological or evolutionary consequences. Studies on variation in plasticity often fail to show IxE or GxE; this can indicate a genuine absence or simply a lack of power. There is, however, another factor that could potentially affect the power to detect IxE or GxE: the choice of the environmental variable included in the analysis. Identifying the genuine environmental driver of phenotypic plasticity will mostly be impossible and hence only a proxy is included in the analysis. However, if this proxy is too weakly correlated with the real driver of plasticity, this will bias IxE and GxE downwards, could lead to spurious negative results and invalid conclusions. As the mean phenotype in a given environment captures all environmental effects on the phenotype, using it as ‘environment’ in the analysis should increase the power to detect IxE or GxE. Using simulations I here showed that using weakly correlated proxies indeed biased estimates downwards but that when using the environment-specific means this was not the case. Using environment-specific means as a covariate has been commonly used in animal and plant breeding but rarely used outside these fields despite its potential usefulness as ‘yardstick’ to test whether IxE or GxE is present or absent in the phenotype of interest.

## Introduction

Many traits are phenotypically plastic (Pigliucci, 2005), i.e. individuals or genotypes express different phenotypes in different environments. Phenotypic plasticity is generally described by a ‘reaction norm’ (Woltereck, 1909), which is defined by its intercept, i.e. the phenotype in the average environment, and its slope, i.e. the ‘sensitivity’ to the environment, in the case of a simple linear reaction norm. Genotypes and individuals can differ in their reaction norm slopes, in which case we talk about genotype-by-environment (GxE) and individual-by-environment interactions (IxE) (e.g. Nussey, Wilson & Brommer, 2007). Understanding whether and how much individual and genetic variation in reaction norms is present in a trait is biologically important. For example, low ‘consistency’, or repeatability, of individuals across different environments could be due to crossing reaction norms, i.e. high IxE variation, or a large environmental component, i.e. high residual variation. Disentangling these two explanations would obviously be important as the biological inferences that would be drawn would be very different. Furthermore, variation in reaction norm slopes could explain a lacking selection response in heritable traits. Even if selection acts in the same direction in all environments, low consistency of individual phenotypes across environments, caused by high IxE and hence crossing reaction norms, can lead to no net selection at the individual level across all environments thereby explaining a lacking selection response (e.g. Turelli & Barton, 2004; Kokko & Heubel, 2008).

Phenotypic plasticity enables populations to cope with changing environments (e.g. Yeh & Price, 2004) but in the long term such plastic responses are unlikely to be sufficient and the reaction themselves will have to evolve (Gienapp, Reed & Visser, 2014). Knowing whether and how much genetic variation in reaction norms is present is then crucial to assess whether populations will be able to cope with environmental change in the long term. Many studies explored individual and genetic variation in reaction norms in behaviour, phenology, i.e. the seasonal timing of life-cycle events, or physiology (e.g. Sih, Bell & Johnson, 2004; Reed et al., 2009; Hau & Goymann, 2015; Stedman, Hallinger, Winkler & Vitousek, 2017). While some studies found statistically significant variation in reaction norms (e.g. Nussey, Postma, Gienapp & Visser, 2005; Brommer, Rattiste & Wilson, 2008; Reed et al., 2009; Stedman et al., 2017), others did not (e.g. Reed et al., 2006; Charmantier et al., 2008). Negative results always raise the question whether it is a genuine absence, in this case of individual or genetic variation in reaction norms, or whether the study had enough power to detect this variation. Two simulation studies showed that the power to detect individual variation in reaction norms depends on both the number of observations per individual and the number of individuals sampled (Martin, Nussey, Wilson & Réale, 2011; van de Pol, 2012) but the choice of the environmental variable in the analysis could also play a role.

The majority of studies on individual and genetic variation used a specific form of the mixed model, the random regression or infinite dimensional model (Henderson, 1982; Kirkpatrick, 1989; Morrissey & Liefting, 2016). Instead of regarding phenotypes at discrete environmental states as separate traits and modelling their variances and covariances, here the trait and the (co)variance in reaction norm slope and intercept is modelled as a continuous (linear) function of the environment. Intercepts and slopes of individual reaction norms are modelled as random effects, i.e. random draws from a normal distribution with mean zero and (co)variances to be estimated. This approach means that a continuous environmental variable is needed as covariate against which the phenotypes are regressed. The choice of this environmental could, however, also affect the power to detect variation in reaction norms, which, so far, has been underappreciated. Using a non-meaningful covariate will lead to a downward bias in the population-level reaction norms and also to a downward bias in individual and genetic variation of reaction norms.

In experimental work it is obvious which variable to choose as covariate as this will be the variable that was experimentally manipulated. In field work, however, a large number of variables could affect the trait and are hence potential candidates to be used as covariate in an analysis of phenotypic plasticity. Biological knowledge will obviously help with the choice of this variable. For example, phenological traits, as, e.g., flowering, migration or breeding, generally depend on ambient temperatures (Sparks & Carey, 1995; Visser, Holleman & Caro, 2009) and hence local temperatures would be an obvious choice as covariate. There are, however, many different ways to measure temperature as it can be averaged or summed over an almost infinite number of periods. This issue arises for virtually any trait in a natural population as there will always be more than one environmental variable to choose from. To complicate matters further traits can, of course, depend on more than one variable. For example, it has been shown that breeding time is affected by spring temperature and population density in tree swallows (*Tachycineta bicolor*) (Bourret, Bélisle, Pelletier & Garant, 2015) or spring temperature in interaction with day length in great tits (*Parus major*) (Gienapp, Hemerik & Visser, 2005). Consequently, it will very rarely be possible to include the real driver of plasticity as covariate in the analysis, simply because it is not known, and instead a proxy that correlates with the real causal driver(s) of plasticity has to be used.

If this proxy is, however, only weakly correlated to the real driver of plasticity, this will lead to an underestimation of population-level plasticity and to an underestimation of variation in reaction norm slopes. This phenomenon can be related to ‘attenuation’ (Spearman, 1904), which he described as the lowered correlation between two variables due to measurement error in them. If we regress phenotypes against a proxy that is not perfectly correlated with the true driver, i.e. *r*<1, and regard the reduced correlation between proxy and true driver as measurement error in the true driver, the observed correlation between proxy and phenotypes will be reduced by a factor inverse to the square root of the correlation between proxy and true driver.

Given sufficient sample size, any absence of individual variation in reaction norm slopes can hence indicate a true absence, i.e. no variation in reaction norm slopes, or that an unsuitable proxy, too weakly correlated with the real driver(s) of plasticity, was used. For example, Brommer *et al*. (2005) analysed IxE in breeding time in collared flycatchers (*Ficedula hypoleuca*) with three different environmental variables, temperature, rainfall, and NAO. All three variables explained significant variation in the trait but statistically significant IxE was found for only two of these variables. Similarly, no statistically significant IxE was found for breeding time in the Wytham Wood great tit (*Parus major*) population when temperature sums from 1 March to 25 April were used as a covariate (Charmantier et al., 2008), while using the mean temperature from 15 February to 25 April as a covariate resulted in statistically significant IxE (Husby et al., 2010).

The main problem in such analyses is that it will be virtually impossible to be sure that the chosen proxy is correlated closely enough with the true driver of plasticity to allow the detection of IxE or GxE. One potential solution to this is regressing individual phenotypes against environment-specific mean phenotypes. This, seemingly circular, method, called ‘joint-regression analysis’ or ‘Finlay-Wilkinson regression’, is commonly used in animal and plant breeding studies. It was first proposed by Yates and Cochran (1938) but only later made popular by Finlay and Wilkinson (Finlay & Wilkinson, 1963). It is widely used in animal and plant breeding to test reliably whether the (relative) merit of genotypes is constant across environments, i.e. whether GxE exists or not (Lynch & Walsh, 1998; James, 2009). Since the mean phenotype in each environment incorporates all plastic effects of environmental drivers, it would be correlated closely with the true, but unknown, driver of plasticity.

I here explored how the correlation of a proxy environmental variable with the real driver of plasticity can bias estimates of individual variation in plasticity and how the ‘Finlay-Wilkinson regression’ approach compares to using these various proxies. To do this, I regressed in a random regression model simulated phenotypes against environment-specific phenotypic means and against nine different environmental variables that are correlated with the real driver with an *r* of 0.9 to 0.1 but did not themselves have an effect on the phenotypes, i.e. are only proxies. In the simulated data, individual variation in reaction norms (IxE) was present and the key point of interest was with which proxy it could be detected.

## Methods-Simulation Model

Individual phenotypes were simulated with an ‘individual-based model’. Individual reaction norm (RN) intercepts and slopes were drawn from normal distributions with means of zero and variances of 5 and 0.1, respectively. Individual intercepts and slopes were assumed to be uncorrelated. The true driver of plasticity, environmental variable E1, was simulated for 20 years (i.e. one value per year for each variable across 20 years) by drawing values from a normal distribution with a mean of zero and a variance of one. Nine other environmental variables (E2 to E10) that were correlated with E1 by 0.9 to 0.1 were simulated using the following equation:

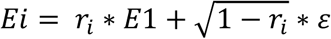

with *E*_*i*_ being the *i*th environmental variable, *r*_*i*_ being its correlation with *E*1 and *ε* being a random error drawn from a normal distribution with mean 0 and variance 1.

Phenotypes of 500 individuals were simulated by first determining a ‘birth year’ that was drawn randomly from the years but ensuring that the four observations of each individual would fit within its ‘life time’. Phenotypes were simulated as follows:

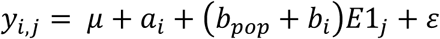

with *y*_*i,j*_ being the phenotype of individual *i* in year *j*, *μ* being the population-level RN intercept, *a*_*i*_ being the individual effect for RN intercept, *b*_*pop*_ being the population-level RN slope, *b*_*i*_ being the the individual effect for RN slope, E1_*j*_ being the value of environmental variable E1 (i.e. the true causal driver of plasticity) in year *j*, and *ε* being a random error with mean zero and variance of 0.25. Additionally, scenarios with different age distributions, roughly resembling a short-lived passerine and a long-lived mammal were explored, resulting in different distributions of number of observations per individual, see Supplementary Material.

To test whether additional but unidentified variables can bias results from an analysis with environment-specific means more than results from analyses with environmental variables, three such variables were modelled: age, habitat effects, and a systematic time trend, akin to a genetic change as response to a constant selection pressure. Age effects were -0.5, 0, 0.5, and 0.3 for ages of 1 to 4, roughly simulating an increase with an optimum at age 3 and weak senescence for age 4. Habitat effects were modelled by randomly distributing individuals over 5 different habitats with habitat-specific effects of -0.1, -0.05, 0.0, 0.05, and 0.1. Individuals could disperse to different habitats throughout their lifetime. The time trend was modelled by letting phenotypes increase by 0.01 per year.

Since the environmental variables varied annually, meaningful environment specific means (ESMs) would be annual means and were hence calculated by averaging the phenotypes over years. Phenotypes were then regressed against all environments and ESMs (all standardised before analysis) in separate analyses using a random regression model corresponding to the model above to estimate individual variation in RN intercepts and slopes. Individual was fitted as a random effect and the environmental variable as a continuous fixed effect. The significance of variation in slopes was tested with a likelihood-ratio test that compared a model with individual variation in intercepts and slopes against a model with only variation in intercepts. Heterogeneous residual variation can lead to spurious individual variation in RN slopes but since all potential variation in phenotypic variation across a given environmental variable would be driven by variation in RN slopes, homogeneous residual variation across the environment was fitted. The difference in model likelihoods is approximately X^2^-distributed with 2 df. For each model with a different environment (E1 to E10 and am) the estimated variances for intercept and slope, the significance for variation in slopes as well as the estimate for the population-level slope were stored. The simulation was repeated 1000 times. Simulation and analysis was done in R 3.1 (R Development Core Team, 2015). The code is provided as Supplementary Material.

## Results-Simulation Model

Unsurprisingly, using the environmental variable that was used to simulate the phenotypes (E1), i.e. the real driver of plasticity, as a covariate in the analysis recovered the input values for reaction norm (RN) slope and individual variation in intercept and slope of 0.5, 5.0, and 0.1, respectively. For the model without additional variables (age, habitat, year) the mean estimate for the population-level slope using E1 was 0.48, while the mean estimated individual variation in intercepts and slope was 5.01 and 0.095, respectively.

The estimates for population-level slope and individual variation in RN slopes clearly declined with a decreasing correlation of the used covariate with the ‘real driver’ (E1) (Fig. 1). As the estimated individual variation in RN slopes declined, the power, i.e. the proportion of replicates, in which IxE was statistically significant, decreased indicating that it would be highly unlikely to detect statistically significant individual variation in RN slopes if the covariate would be correlated with the real driver by 0.5 and less (Fig. 2). The estimates for individual variation in RN slopes from the model using environment-specific means as covariate (mean: 0.092) were as close or closer to the estimates from the model using E1 than models using other environmental variables (E2 to E10) (Fig. 1). Other factors that affect the phenotype but are unaccounted for will alter the environment-specific mean phenotypes and can thereby potentially affect the results. Here, age, a second environmental variable, and a systematic time trend were tested, see Methods, but none of them led to any systematic bias in the probability to detect IxE and its estimates, see Supplementary Material, Figs S1 & 2.

**Fig. 1.**
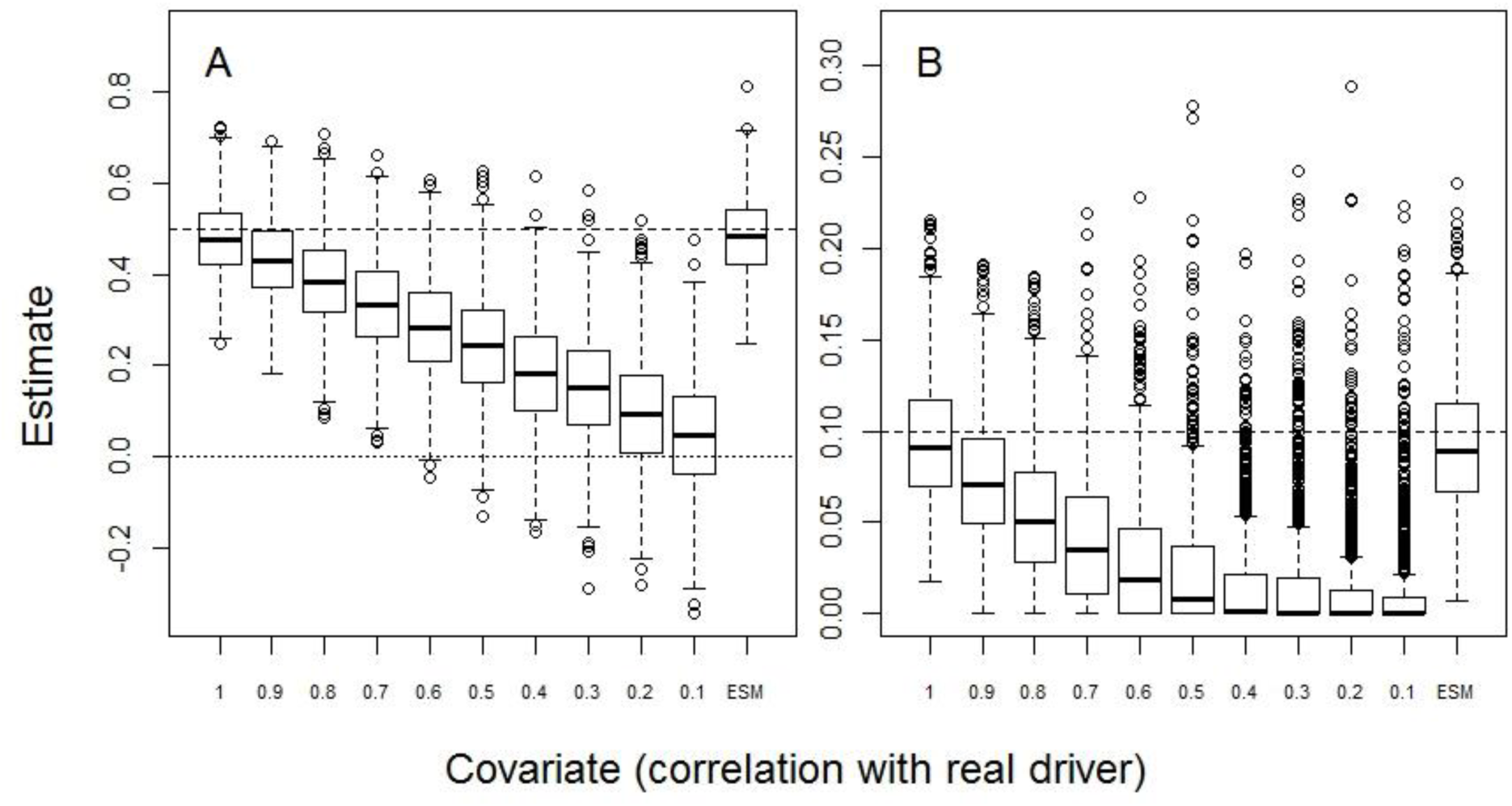
Boxplot of estimates for the population-level reaction norm slope (A) and individual variation in reaction norm slopes (B) depending on the covariate included in the random regression model. The covariates included in the model are indicated by their correlation with the ‘real driver’ of plasticity (E1). ESM indicates environment-specific mean phenotypes as covariate. The dashed lines indicate the input values for the slope and variation in slopes. Bold line indicates median, box margins 1st and 3rd quartile and whiskers 1.5*inter-quartile range.

**Fig. 2.**
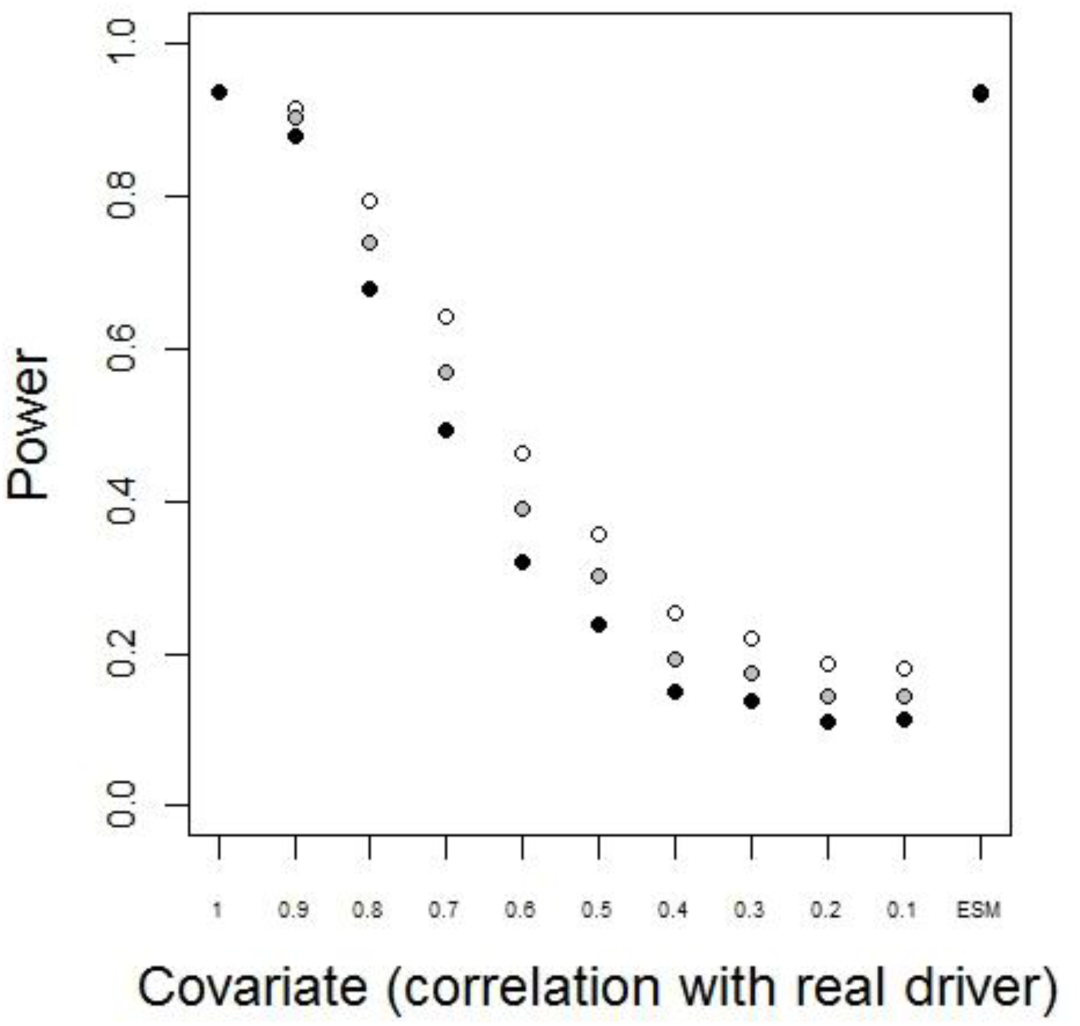
The power to detect variation in reaction norms slopes (IxE) depends on the covariate included in the model. The proportion of replicates, in which IxE was statistically significant at a significance level of 0.05 (open symbols), of 0.01 (grey symbols), and 0.001 (black symbols), is plotted against the correlation of the covariate with the ‘real driver’ of plasticity (E1). ESM indicates environment-specific mean phenotypes as covariate.

Varying the number of observations per individual, following roughly the distributions in a large long-lived mammal, as red deer, and a small, short-lived passerine bird, as the great tit, did not have a large effect on the power to detect IxE in the simulations and did not alter the performance of environment-specific means as covariate, see Supplementary Material, Figs. 3-6.

## Results-Mathematical Derivation

I here derive the correlation between the true driver of plasticity (E1) and the environment-specific mean (ESM). The ESM is the average phenotype in any given environment:

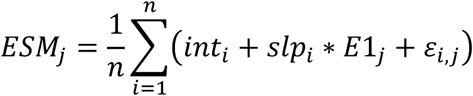

with *int*_*i*_ and *slp*_*i*_ being the intercept and the slope of the reaction norm of individual *i*, *E*1_*j*_ the ‘real driver’ in environment *j*, *ε*_*i,j*_ a random error term, and *n* sample size. *int*_*i*_ and *slp*_*i*_ are random deviations from the population mean values and for infinite sample sizes the averages over all individual intercepts and slopes, 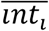 and 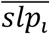, become equal to the population mean values and hence the average of the ESM in any environment *E*1 simply would equal the expectation

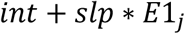

with *int* and *slp* being the population mean values of the reaction norm intercept and slope. However, for limited sample sizes the individual deviations from the population reaction norm plus the additional error term imply that the average becomes

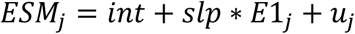

with *u*_*j*_ being the ‘sampling error’ incorporating individual deviations from the population reaction norm and summed individual error terms. This contribution to *ESM*_*j*_ will tend to decrease with sample size *n*.

Now we can write the correlation between E1 and ESM across environments *j* as

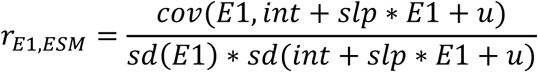

which can be simplified to

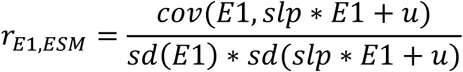
()
since *cov*(*aX* + *bY*, *cW* + *dV*) = *ac cov*(*X*, *W*) + *ad cov*(*X*, *V*) + *bc cov*(*Y*, *W*) + *bd cov*(*Y*, *V*) and *var*(*aX* + *bY*) = *a*^2^*var*(*X*) + *b*^2^*var*(*Y*) + 2*abcov*(*X*, *Y*), the above equation can be re-written as:

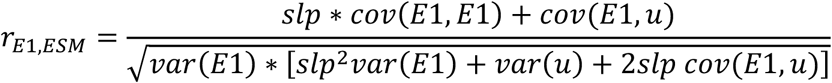

Assuming unbiased sampling such that *cov*(*E*1, *u*) equals zero and using *cov*(*E*1, *E*1) = *var*(*E*1), this can be simplified to:

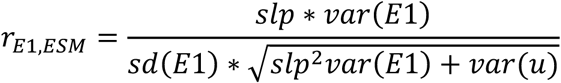

and further to:

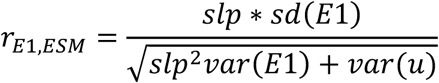

If we now express the ratio of *var*(*u*) and *var*(*E*1) as *x*, then we can re-write the above equation as:

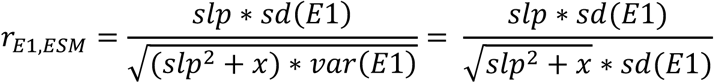

which can be simplified to:

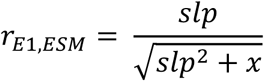

With decreasing *var*(*u*), for example, due to increasing sample size *n*, *x* will approach zero and the above correlation will approach a value of one for positive values of slope *slp* which means that the *ESM* would then outperform any other proxy.

## Discussion

As has been found previously (Brommer et al., 2005; Husby et al., 2010) and systematically explored here, the choice of the environmental variable that affects the phenotype in the reaction norm can affect the probability to detect variation in reaction norm slopes. The less related the included environmental variable is to the real driver of plasticity, the lower the probability to detect statistically significant IxE (Fig. 2). As it is in practice impossible to know how closely the used covariate is correlated to the real driver, it is not useful trying to develop any rules, even rules of thumb, for the power of such analyses. Instead, using environment-specific population means as covariate, i.e. using a so-called ‘Finlay-Wilkinson’ regression (Yates & Cochran, 1938; Finlay & Wilkinson, 1963), could provide an alternative means to explore IxE without the need to identify an environmental variable closely related to the real driver. While this approach does not give biological insight into the environmental variable underlying phenotypic plasticity and IxE in the analysed trait, it can reliably, given sufficient sample size, indicate the presence or absence of IxE.

It could be shown that the correlation between the true driver of plasticity and the environment-specific mean approaches one with decreasing ‘sampling variation’ in the environment-specific mean. This sampling variation will obviously decrease with increasing sample size. However, since it also incorporates variation due to individual (or genetic) variation in reaction norms, it will also decrease with a decrease in variation in reaction norms. This means the smaller the effect of interest becomes, the more precise the estimated environment-specific means become and the better their suitability to detect variation in reaction norms.

If the presence of IxE has been shown using a ‘Finlay-Wilkinson’ regression, further analyses can aim to identify to meaningful biotic or abiotic environmental variables that drive IxE in the trait of interest. If IxE can be found using an environmental variable, it can then be concluded that this variable is reasonably closely correlated with the environmental variable that causally affects the trait. In this sense a ‘Finlay-Wilkinson’ regression can serve as a ‘yardstick’ against that models including other environmental variables can be compared.

Whether or not IxE, and also genetic variation in reaction norms slopes (GxE), is present or not in a population or species, is biologically relevant. For example, IxE can increase or decrease the amount of among-individual variation in novel environments, which affects the opportunity of selection and thereby potential evolutionary change, even in the absence of GxE. The absence of IxE can also be biologically interesting; for example, Reed et al. (2006) found no IxE in breeding time of common guillemots (*Uria aalge*) in response to the North Atlantic Oscillation and argued that this lack of IxE was a consequence of the benefits of coordinated breeding in colonial birds, such as the common guillemot. However, in such cases it would be desirable to be sure that this absence of IxE is not caused by an unsuitable choice of the environmental variable affecting the trait. In fact, when re-analysing this data using phenotypic environment-specific means, in this case annual mean egg-laying dates, as environmental variable, statistically significant IxE was found (T.E. Reed pers. comm.).

Ramakers et al. (2018) recently used environment-specific means as covariate to enable their ‘meta-analysis’ of GxE. To explore whether a coupling of heritability with the strength of selection, caused by GxE and covariation of selection with the environment (SxE), they used environment-specific means in their analysis to ensure that potentially existing GxE was not masked by a badly chosen environmental variable and also because many of the studies re-analysed here did not provide any environmental variable.

Trait expression is obviously not only affected by environmental variables but also by individual ‘state’ variables, such as age or physical condition and potentially other environmental variables, for example habitat in addition to temperature. Not accounting for such variables can bias environment-specific mean phenotypes, for example, if age structure in the population varies from year to year and this could potentially lead to specific biases in estimates from ‘Finlay-Wilkinson’ regressions. This potential bias may be especially problematic when there is a consistent time trend in phenotypes, as caused, for example, by a genetic selection response. Such potential biases would also affect the phenotypes, albeit not the covariate, of analyses based on environmental variables. The results here show that results from ‘Finlay-Wilkinson’ regressions are not biased compared to using environmental variables (see Appendix). It is, however, always preferable to include and test all variables potentially affecting the trait to obtain more accurate estimates of variations in reaction norms and also to gain a better understanding of which abiotic and biotic variables affect the trait.

An important assumption underlying the ‘Finlay-Wilkinson’ regression is that the response to the ‘true’ driver of plasticity is (approximately) linear as strongly non-linear reaction norms, for example, quadratic or sigmoidal, could lead to identical mean phenotypes in different environments, which would obviously interfere with reliably estimating IxE. It is also important that the environment-specific means are based on adequate sample sizes as it has been shown that unreliably estimated environment-specific means can bias estimates of GxE (Calus, Bijma & Veerkamp, 2004) and the same will apply for IxE, too.

Genetic variation in reaction norm slopes (GxE) is also relevant as environmental change likely leads to selection on both reaction norm intercept and slope (Gienapp et al., 2014). How the power to detect GxE depends on the choice of the covariate included in the analysis was not addressed here because of the specific issues with the power of quantitative genetic analyses. The sampling variation in heritability estimates, i.e. the power of quantitative genetic analyses, depends on the sample size but also on the variation of relatedness within the population (Visscher & Goddard, 2015). The variation in relatedness depends on a number of species- or population-specific ecological parameters, as, for example, dispersal or mating system. It would have been possible to simulate GxE but the obtained results would have been difficult to generalise. However, IxE is generally regarded as an ‘upper limit’ for GxE. Having found no IxE with sufficient sample size and using a ‘Finlay-Wilkinson’ regression it would hence be highly unlikely to find GxE in the same population. It should, however, be noted that this ‘yardstick’ cannot universally be applied as in short-lived species necessary sample sizes, in terms of repeat observations of individuals, may not ever be reached. Hence, IxE may not be well estimable but – given suitable relatedness information – quantitative genetic analyses could be possible and significant GxE could be found.

In animal and plant breeding the ‘Finlay-Wilkinson’ regression has long been used but very rarely outside this field (James, 2009). I argued here that it can be useful as a ‘yardstick’ in analyses exploring individual (and genetic) variation in reaction norm slopes as its results are unbiased by the correlation between it and the environmental variable causally affecting the trait. This can be especially relevant for studies not finding statistically significant IxE and could therefore give us a better understanding how prevalent (or not) individual-by-environment interactions really are.

## Supporting information

## Acknowledgements

M. E. Visser, T. E. Reed, H. A. Mulder and T. Van Dooren commented on the manuscript and provided useful feedback. T. E. Reed kindly re-analysed the isle of May data on guillemot breeding and allowed me to cite the results here. Two anonymous reviewers provided constructive criticism.

## References

Bourret, A., Bélisle, M., Pelletier, F., Garant, D. (2015) Multidimensional environmental influences on timing of breeding in a tree swallow population facing climate change. Evolutionary Applications 8: 933–944.

Brommer, J. E., Merilä, J., Sheldon, B. C., Gustafsson, L. (2005) Natural selection and genetic variation for reproductive reaction norms in a wild bird population. Evolution 59: 1362–1372.

Brommer, J. E., Rattiste, K., Wilson, A. J. (2008) Exploring plasticity in the wild: laying date-temperature reaction norms in the common gull *Larus canus*. Proceedings of the Royal Society B: Biological Science 275: 687–693.

Calus, M. P., Bijma, P., Veerkamp, R. F. (2004) Effects of data structure on the estimation of covariance functions to describe genotype by environment interactions in a reaction norm model. Genetics Selection Evolution 36: 489.

Charmantier, A., McCleery, R. H., Cole, L. R., Perrins, C., Kruuk, L. E. B., Sheldon, B. C. (2008) Adaptive phenotypic plasticity in response to climate change in a wild bird population. Science 320: 800–803.

Finlay, K. W., Wilkinson, G. N. (1963) The analysis of adaptation in a plant-breeding programme. Australian Journal of Agricultural Research 14: 742–754.

Gienapp, P., Hemerik, L., Visser, M. E. (2005) A new statistical tool to predict phenology under climate change scenarios. Global Change Biology 11: 600–606.

Gienapp, P., Reed, T. E., Visser, M. E. (2014) Why climate change will invariably lead to selection on phenology. Proceedings of the Royal Society B-Biological Sciences 281: 20141611.

Hau, M., Goymann, W. (2015) Endocrine mechanisms, behavioral phenotypes and plasticity: known relationships and open questions. Frontiers in Zoology 12.

Henderson, C. R. (1982) Analysis of covariance in the mixed model: higher-level, non-homogeneous, and random regressions. Biometrics 38: 623–640.

Husby, A., Nussey, D. H., Visser, M. E., Wilson, A. J., Sheldon, B. C., Kruuk, L. E. B. (2010) Contrasting patterns pf phenotypic plasticity in reproductive traits in two great tit (Parus major) populations. Evolution 64: 2221–2237.

James, J. W. (2009) Genotype by envionment interaction in farm animals. In: van der Werf, J., Graser, H.-U., Frankham, R. and Gondro, C., eds. Adaptation and fitness in animal populations: Evolutionary and breeding perspectives on genetic resource management. 151–167. Springer.

Kirkpatrick, M. (1989) A quantitative genetic model for growth, shape, reaction norms, and other infinite-dimensional characters. Journal of Mathematical Biology 27: 429–450.

Kokko, H., Heubel, K. (2008) Condition-dependence, genotype-by-environment interactions and the lek paradox. Genetica 134: 55–62.

Lynch, M., Walsh, B. (1998) Genetics and analysis of quantitative traits. Sunderland, Massachusetts: Sinauer.

Martin, J. G. A., Nussey, D. H., Wilson, A. J., Réale, D. (2011) Measuring individual differences in reaction norms in field and experimental studies: a power analysis of random regression models. Methods in Ecology and Evolution 2: 362–374.

Morrissey, M. B., Liefting, M. (2016) Variation in reaction norms: Statistical considerations and biological interpretation. Evolution 70: 1944–1959.

Nussey, D. H., Postma, E., Gienapp, P., Visser, M. E. (2005) Selection on heritable phenotypic plasticity in a wild bird population. Science 310: 304–306.

Nussey, D. H., Wilson, A. J., Brommer, J. E. (2007) The evolutionary ecology of individual plasticity in wild populations. Journal of Evolutionary Biology 20: 831–844.

Pigliucci, M. (2005) Evolution of phenotypic plasticity: where are we going now? Trends in Ecology & Evolution 20: 481–486.

R Development Core Team (2015) R: A language and environment for statistical computing. Vienna, Austria: R Foundation for Statistical Computing.

Ramakers, J. J. C., Culina, A., Visser, M. E., Gienapp, P. (2018) Environmental coupling of heritability and selection is rare and of minor evolutionary significance in wild populations. Nature Ecology & Evolution 2: 1093–1103.

Reed, T. E., Wanless, S., Harris, M. P., Frederiksen, M., Kruuk, L. E. B., Cunningham, E. J. A. (2006) Responding to environmental change: plastic responses vary little in a synchronous breeder. Proceedings of the Royal Society B 273: 2713–2719.

Reed, T. E., Warzybok, P., Wilson, A. J., Bradley, R. W., Wanless, S., Sydeman, W. J. (2009) Timing is everything: flexible phenology and shifting selection in a colonial seabird. Journal of Animal Ecology 78: 376–387.

Sih, A., Bell, A., Johnson, J. C. (2004) Behavioral syndromes: an ecological and evolutionary overview. Trends in Ecology & Evolution 19.

Sparks, T. H., Carey, P. D. (1995) The Responses of Species to Climate over 2 Centuries - an Analysis of the Marsham Phenological Record, 1736-1947. Journal of Ecology 83: 321–329.

Spearman, C. (1904) The proof and measurement of association between two things. American Journal of Psychology 15: 72–101.

Stedman, J. M., Hallinger, K. K., Winkler, D. W., Vitousek, M. N. (2017) Heritable variation in circulating glucocorticoids and endocrine flexibility in a free-living songbird. Journal of Evolutionary Biology 30: 1724–1735.

Turelli, M., Barton, N. H. (2004) Polygenic variation maintained by balancing selection: Pleiotropy, sex-dependent allelic effects and GxE interactions. Genetics 166: 1053–1079.

van de Pol, M. (2012) Quantifying individual variation in reaction norms: how study design affects the accuracy, precision and power of random regression models. Methods in Ecology and Evolution 3: 268–280.

Visscher, P. M., Goddard, M. E. (2015) A general unified framework to assess the sampling variance of heritability estimates using pedigree or marker-based relationships. Genetics 199: 223–232.

Visser, M. E., Holleman, L. J. M., Caro, S. P. (2009) Temperature has a causal effect on avian timing of reproduction. Proceedings of the Royal Society B 276: 2323–2331.

Woltereck, R. (1909) Weitere experimentelle Untersuchungen über Artveränderung, speziel über das Wesen quantitativer Artunterschiede bei Daphniden. Verhandlungen der Deutschen Zoologischen Gesellschaft 19: 110–173.

Yates, F., Cochran, W. G. (1938) The analysis of groups of experiments. Journal of Agricultural Science 28: 556–580.

Yeh, P. J., Price, T. D. (2004) Adaptive phenotypic plasticity and the successful colonization of a novel environment. American Naturalist 164: 531–542.

