## Supplementary material for "The choice of the environmental covariate affects the power to detect variation in reaction norm slopes"

**Supplementary Figures**

To test whether additional but unidentified variables can bias results from an analysis with environment-specific means more than results from analyses with environmental variables, three such variables were modelled: age, habitat effects, and a systematic time trend, akin to a genetic change as response to a constant selection pressure. As can be seen from the figures below neither age (Fig.S1), habitat effects (Fig.S2), nor a systematic time trend (Fig.S3) lead to bias in the estimates using environment-specific means as covariate. The power of analyses including these variables was also unaffected, see Fig.S4.

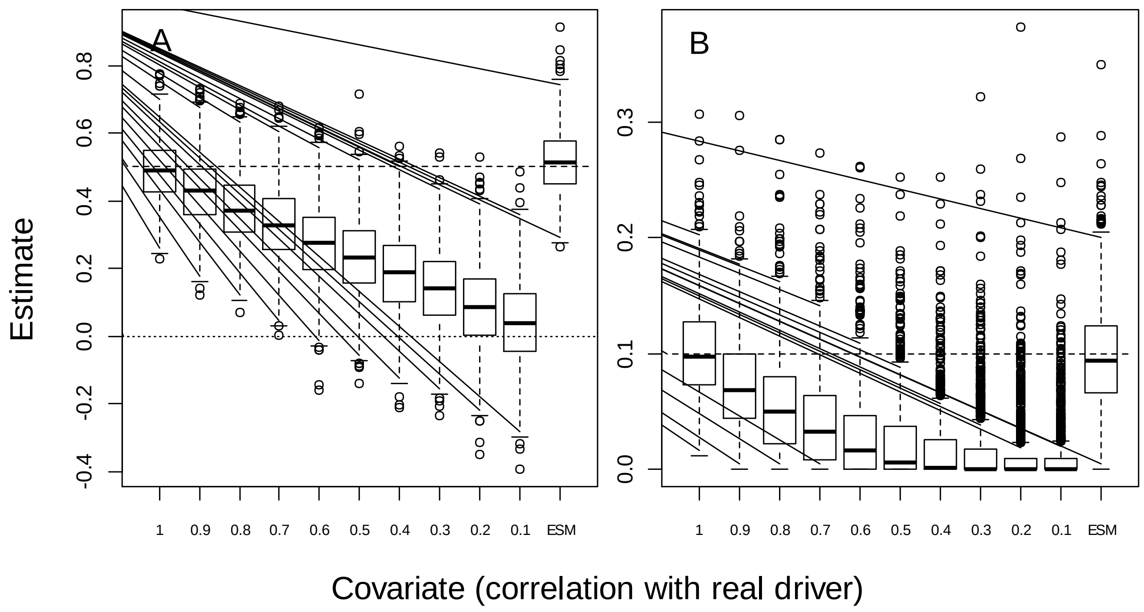


Fig. S1. Boxplot of estimates for the population-level reaction norm slope (A) and individual variation in reaction norm slopes (B) depending on the covariate included in the random regression model when the model included an age effect that was unaccounted for. The covariates included in the model are indicated by their correlation with the ‘real driver’ of plasticity (E1). ESM indicates environment-specific mean phenotypes as covariate. The dashed lines indicate the input values for the slope and variation in slopes. Bold line indicates median, box margins 1st and 3rd quartile and whiskers 1.5*inter-quartile range.

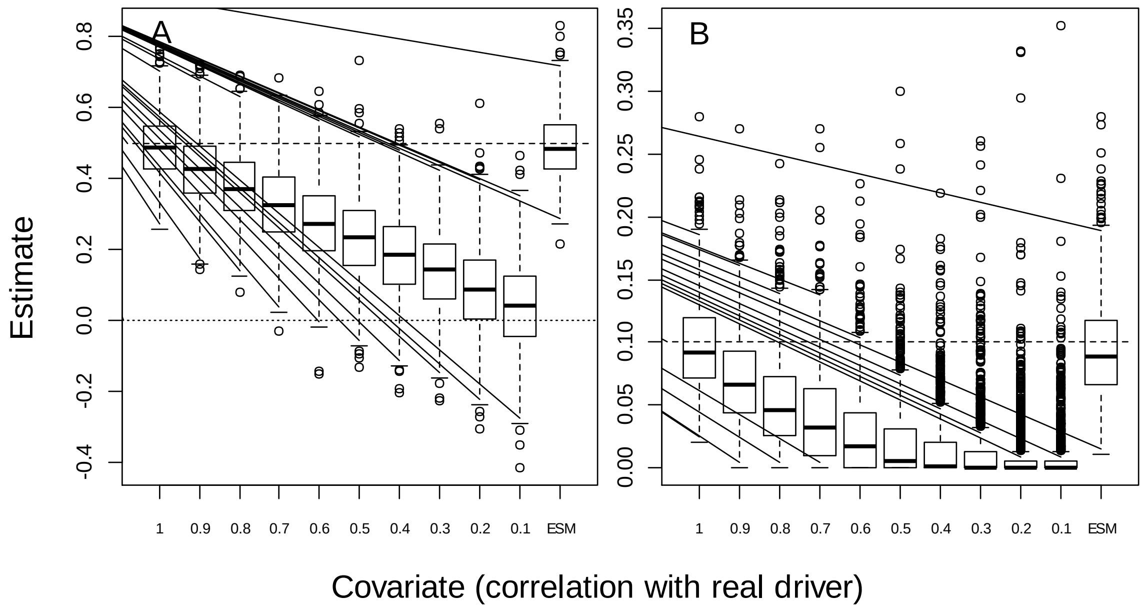


Fig. S2. Boxplot of estimates for the population-level reaction norm slope (A) and individual variation in reaction norm slopes (B) depending on the covariate included in the random regression model when the model included a habitat effect that was unaccounted for.

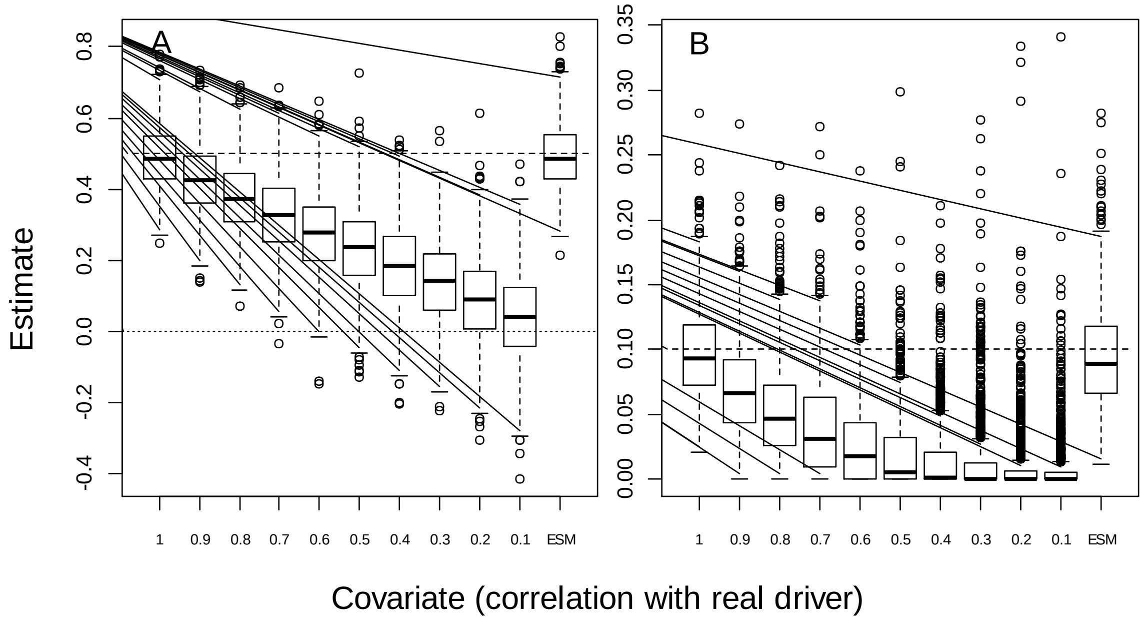


Fig. S3. Boxplot of estimates for the population-level reaction norm slope (A) and individual variation in reaction norm slopes (B) depending on the covariate included in the random regression model when the model included a habitat effect that was unaccounted for.

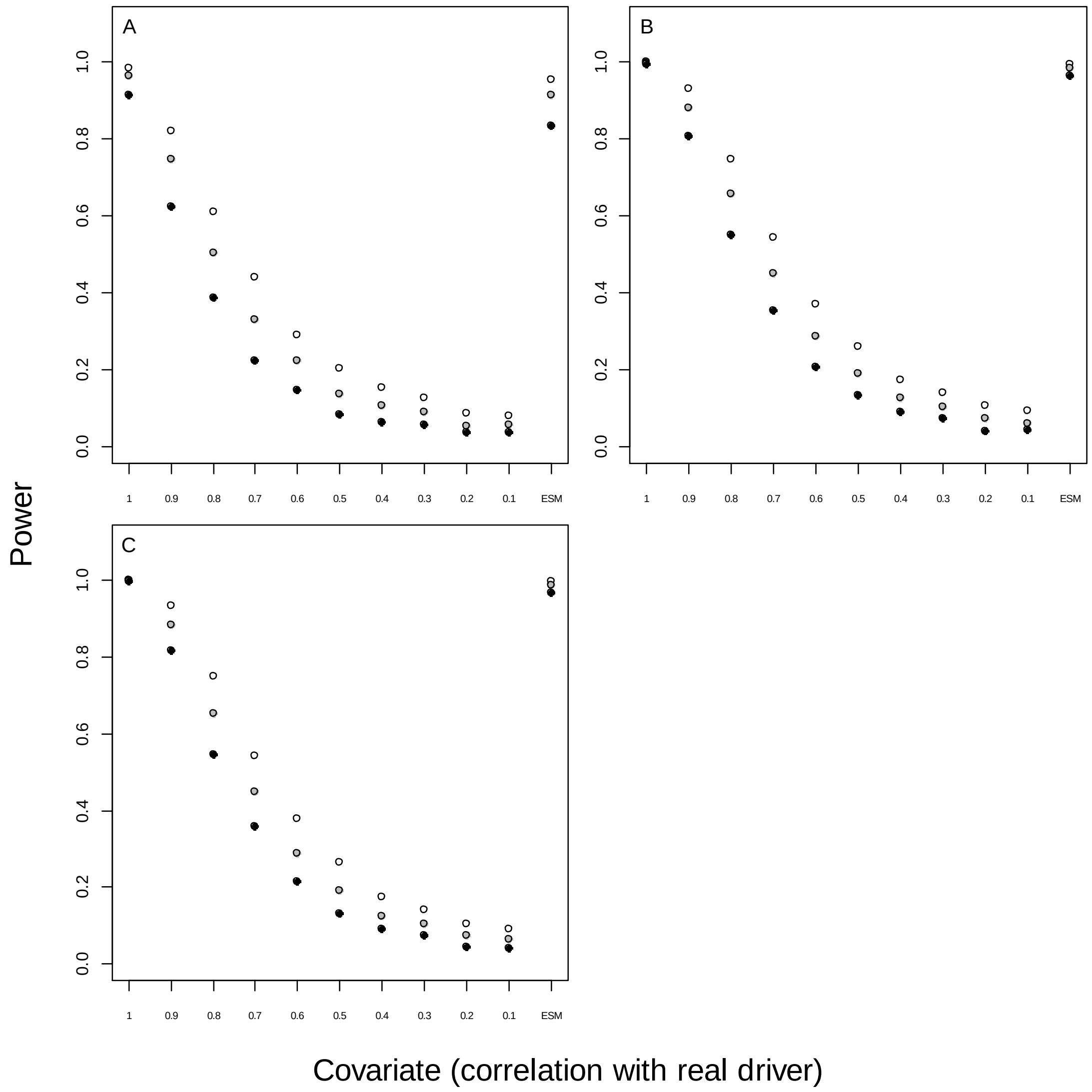


Fig. S4. The power to detect variation in reaction norms slopes (IxE) in models including age (A), habitat (B), and a systematic time trend (C) as un-accounted for variables depending on the covariate included in the model. The proportion of replicates, in which IxE was statistically significant at a significance level of 0.05 (open symbols), of 0.01 (grey symbols), and 0.001 (black symbols), is plotted against the correlation of the covariate with the ‘real driver’ of plasticity (E1). ESM indicates environment-specific mean phenotypes as covariate.
